## Supplementary Figure for "A nanobody-based fluorescent reporter reveals human α-synuclein in the cell cytosol"

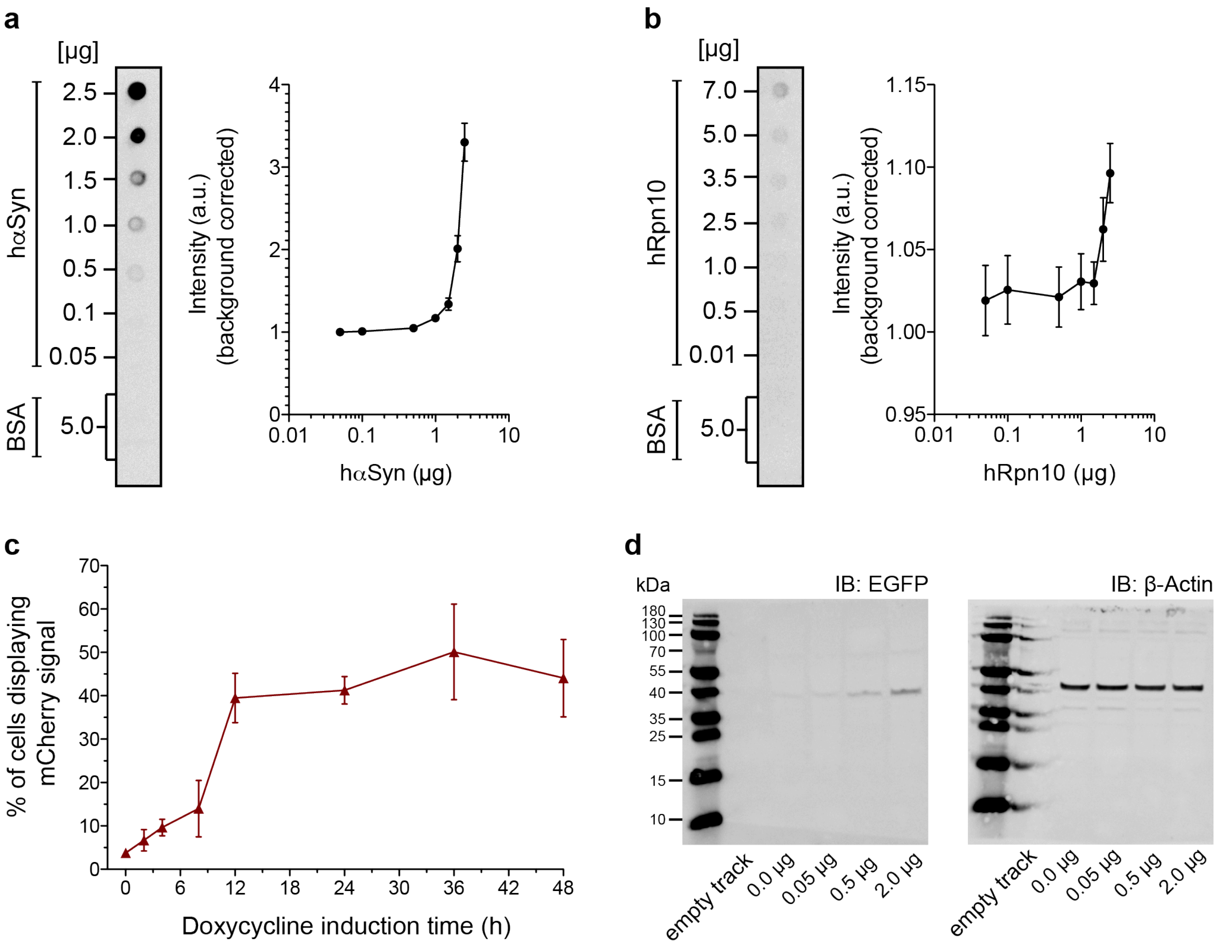


**Supp. Figure 1| NbSyn87 binding to hαSyn and hRpn10 and the biochemical characterization of Reporter-cells**. (**a, b**) Increasing amounts of hαSyn (**a**) or hRpn10 (**b**) were spotted on a nitrocellulose membrane and detected using NbSyn87 directly coupled to Alexa 647. BSA was spotted as control protein and its signal was used as background for normalization. Error bars represent the s.e.m from 3 independent experiments. (**c**) Induction response curve of the Reporter-cells to determine the optimal duration of doxycycline administration at a concentration of 0.5 µg/ml. Error bars represent the s.e.m from 4 independent experiments. (**d**) Full Western blots membranes from Fig. 1f.

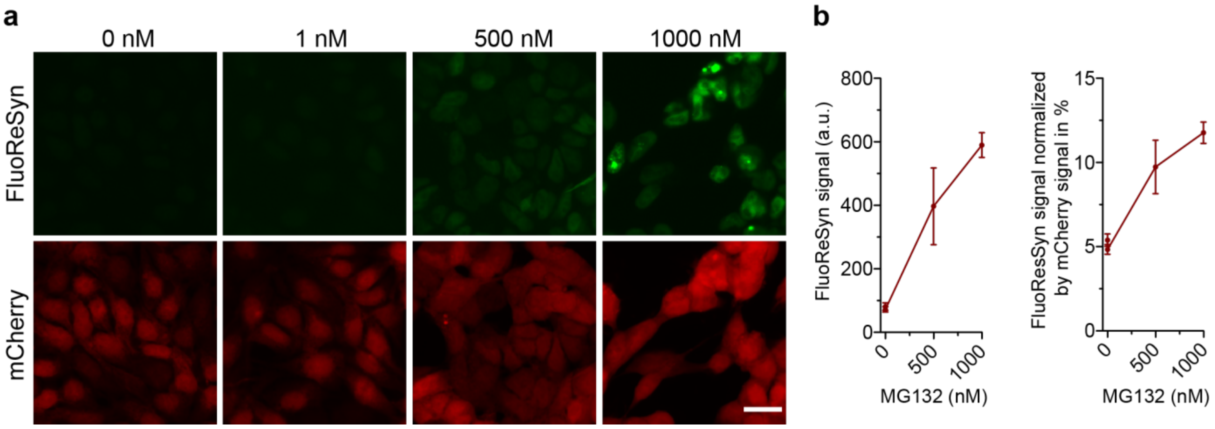


**Supp. Figure 2 | Determination of the optimal concentration for MG132 administration.** (**a**) Fully induced Reporter-cells were exposed to different concentrations of MG132 for 16h. The mCherry signal (red) indicate that the cells are producing FluoReSyn, however, only starting from 500 nM of MG132, the FluoReSyn signal begins to appear (green). (**b**) Quantification of the FluoReSyn signal (green) in arbitrary units (a.u.) or normalized to the signal of mCherry. Error bars represent the s.e.m from 3 independent experiments with several hundreds of cells analyzed per experiment.

**a**

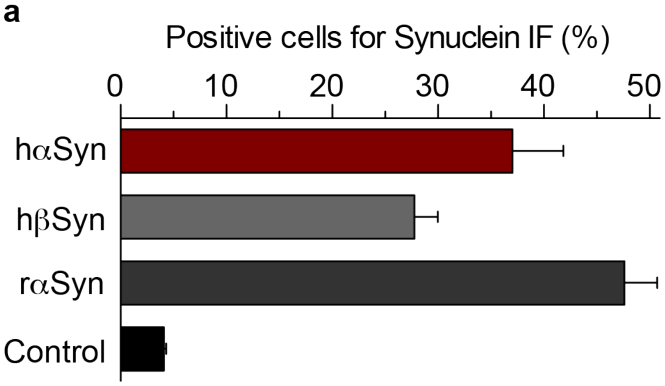


**b**

NbSyn87 putative epitope

```
human αSyn (NP_000336)          111 GILEDMPVDPDNEAYEMPSEEGYQDYEPEA 140
mouse αSyn (NP_001035916.1)     111 GILEDMPVDPGSEAYEMPSEEGYQDYEPEA 140
rat αSyn (NP_062042.1)          111 GILEDMPVDPSSEAYEMPSEEGYQDYEPEA 140
human βSyn (NP_001001502.1)     104 EPLIEPLMEPEGESYEDPPQEEYQEYEPEA 134
```

**Supp. Figure 3 | Specificity of FluoReSyn for hαSyn.** (**a**) Quantification of the signal obtained after immunostaining synucleins with a pan-Synuclein antibody. Error bars represent the s.e.m from 3 independent experiments. (**b**) Amino acid sequence alignment of the putative epitope recognized by NbSyn87 for hαSyn (letters in light purple), mouse and rat αSyn or human βSyn. (accession numbers for each sequence are on the figure). Red letters show a match amino acid to the putative epitope, black letters represent mismatches. The core difference between the sequences is highlighted by a background box.

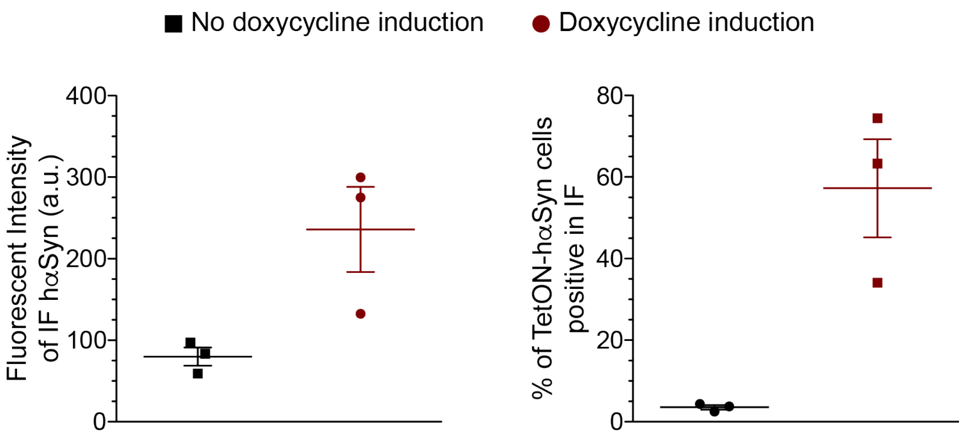


**Supp. Figure 4 | TetON induction of stably-transfected HEK293 cells expressing untagged hαSyn.** (left plot) Fluorescence intensity of cells immunostained for synuclein after being induced or not with 0.5 µg/ml of doxycycline for 16h. (right plot) Percentage of cells displaying a positive immunofluorescense signal for synuclein. Error bars represent the s.e.m from 3 independent experiments.

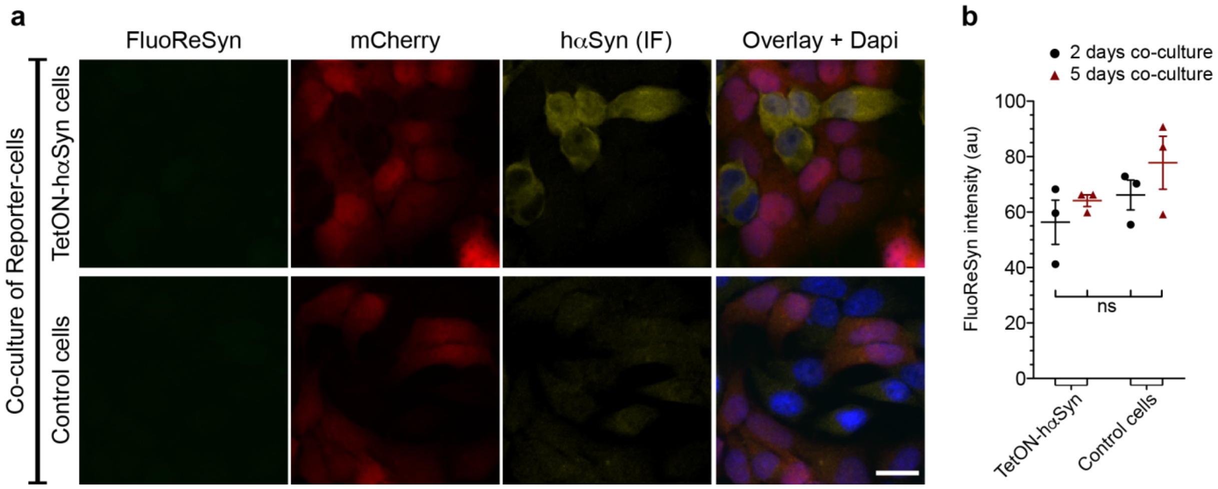


**Supp. Figure 5 | Co-culture of Reporter cells with TetON-hαSyn cells.** (**a**) Reporter-cells co-cultured with either a doxycycline inducible hαSyn expressing cell line (TetON-hαSyn) or the wildtype HEK293 cell line without endogenous hαSyn expression. Induced Reporter-cells can be identified by their mCherry signal while induced TetON-hαSyn cells can be identified by the hαSyn immunofluorescence (IF, yellow). (**b**) Quantitative analysis of FluoReSyn signal intensity of cells with mCherry above 300 AU after 2 or 5 days of co-culturing. Scatter plots show 3 independent experiments ± sem for all condition. Significance was assessed by One-Way ANOVA and Tukey's Post-hoc test. ns, non-significant. Per replication and condition more than 550 cells were analyzed.

**Supp. Table 1**. Demographic and clinical characteristics of the individuals in the CSF study

| ID | Gender | Age | Diagnosis |
|---|---|---|---|
| 1 | f | 64 | Muscular pain-fasciculation syndrome |
| 2 | f | 73 | RLS |
| 3 | m | 77 | FTD |
| 4 | f | 60 | RLS |
| 5 | m | 73 | CBD |
| 6 | m | 74 | SCA |
| 7 | m | 68 | PSP |
| 8 | f | 82 | PSP |
| 9 | f | 71 | PSP |
| 10 | f | 64 | FTD |
| 11 | m | 87 | Vascular parkinsonism |
| 12 | m | 88 | Essential tremor |
| 13 | m | 76 | Vascular dementia |
| 14 | f | 81 | PSP |
| 15 | m | 70 | PSP |
| 16 | f | 81 | NPH without evidence for other neurodegenerative disorders |
| 17 | f | 73 | CBD |
| 18 | m | 76 | RLS |
| 19 | m | 73 | NPH without evidence for other neurodegenerative disorders |
| 20 | f | 69 | NPH without evidence for other neurodegenerative disorders |
| 21 | f | 77 | NPH without evidence for other neurodegenerative disorders |
| 22 | m | 75 | PSP |
| 23 | f | 73 | CBD |
| 24 | f | 64 | SCA |
| 25 | m | 60 | PSP |
| 26 | f | 75 | SCA |
| 27 | f | 77 | Steroid responsive encephalopathy associated with autoimmune thyroiditis |
| 28 | f | 71 | Essential tremor |
| 29 | m | 69 | PSP |
| 30 | f | 75 | Dystonic tremor |
| 31 | m | 84 | PNP |
| 32 | f | 35 | PNP |
| 33 | f | 65 | Neuroleptic-induced dyskinesia |
| 34 | m | 48 | RLS |
| 35 | f | 59 | PSP |
| 36 | m | 76 | PNP |
| 37 | m | 65 | NPH without evidence for other neurodegenerative disorders |
| 38 | m | 60 | SCA |
| 39 | f | 71 | CBD |
| 40 | f | 80 | Benign paroxysmal positional vertigo |
| 41 | f | 73 | RLS |
| 42 | m | 67 | PSP |

m, male; f, female; CBD, corticobasal dementia; FTD, frontotemporal dementia; NPH, normal pressure hydrocephalus; PNP, peripheral neuropathy; PSP, progressive supranuclear palsy; RLS, restless legs syndrome; SCA, spinocerebellar ataxia.
